# Antigen and G-Protein Coupled Receptor signaling differentially control CD8 T cell motility immediately before and after virus clearance in a primary infection

**DOI:** 10.1101/2019.12.16.877845

**Authors:** Kris Lambert-Emo, Emma C. Reilly, Michael Sportiello, David J. Topham

## Abstract

In mice, experimental influenza virus infection stimulates CD8 T cell infiltration of the airways. Virus is cleared by day 9, and between days 8 and 9 there is an abrupt change in CD8 T cell motility behavior transitioning from low velocity and high confinement on day 8, to high velocity with continued confinement on day 9. We hypothesized that it is loss of virus and/or antigen signals in the context of high chemokine levels that drives the T cells into a rapid surveillance mode. Virus infection induces chemokine production, which may change when the virus is cleared. We therefore sought to examine this period of rapid changes to the T cell environment in the tissue and seek evidence on the roles of peptide-MHC and chemokine receptor interactions. Experiments were performed to block G protein coupled receptor (GPCR) signaling with Pertussis toxin (Ptx). Ptx treatment generally reduced cell velocities and mildly increased confinement, except on day 8 when velocity increased and confinement was relieved, suggesting chemokine mediated arrest. Blocking specific peptide-MHC with monoclonal antibody unexpectedly decreased velocities on days 7 through 9, suggesting TCR/peptide-MHC interactions promote cell mobility in the tissue. Together, these results suggest the T cells are engaged with antigen bearing and chemokine producing cells that affect motility in ways that vary with the day after infection. The increase in velocities on day 9 were reversed by addition of specific peptide, consistent with the idea that antigen signals become limiting on day 9 compared to earlier time points. Thus, antigen and chemokine signals act to alternately promote and restrict CD8 T cell motility until the point of virus clearance, suggesting the switch in motility behavior on day 9 may be due to a combination of limiting antigen in the presence of high chemokine signals as the virus is cleared.

## Introduction

Influenza viruses infect roughly 12 percent of the population in any given year [1]. This leads to lost productivity, hospitalizations, and deaths. In the 2017-18 season there was a record 80,000 deaths in the US alone [2]. In 2018-19, the northern hemisphere experienced the longest flu season in over 20 years [3]. Understanding how the immune system controls influenza infection is paramount to the development of improved vaccination strategies and for understanding the disease process itself. Cytotoxic CD8 T cells are responsible for the initial clearance of infected cells, especially in a primary infection when there are no pre-existing antibodies or other types of adaptive immunity [4, 5]. In order to reach the site of infection, the trachea and airway epithelium, the CD8 T cells must traffic through the circulation, exit into the tissue, and migrate within the tissue before crossing into the epithelial surface. There are many things in the tissue microenvironment that the T cells must interact and communicate with, and many features change over the course of an infection as the immune response progresses and the virus gets cleared, which is between day 8 and 9 of the infection.

In the mouse model of influenza infection, virus replication peaks 3-5 days after inoculation [6, 7]. CD8 T cells appear in the tissue beginning around 5-6 days, after which virus titers begin to decrease, and T cell numbers peak at day 8 [5, 8]. As the virus is cleared between day 8 and 9, there is a logarithmic drop in the number of T cells in the lung and airways. Presumably, the end of the infection produces a change in signals that recruit or retain the T cells. It is believed that most of the virus specific T cells die by apoptosis, though it’s unclear if this happens in the tissue or after the T cells leave the tissue, and may be a combination of both.

Our lab developed a model of influenza tracheitis that we use to perform imaging of immune cell motility by intravital multiphoton microscopy (IVMPM) [9]. IVMPM can optically penetrate the entire depth of the trachea once it is exposed by minor surgery [9, 10]. Using genetically engineered CD8 T cells that are fluorescent and recognize an ovalbumin (OVA) peptide presented by H2 Kb class I major histocompatibility complex (MHC) proteins (OT-I-GFP CD8 T cells) [11, 12], and a genetically modified influenza virus that expresses the OVA peptide in the hemagglutinin of the virus [13], we can image the pseudo-virus-specific OT-I T cells as they migrate in the infected trachea. As CD8 T cells infiltrate the tissue, they progressively accumulate and gradually become arrested and confined by day 8. We previously reported that there is an abrupt change in motility behavior between day 8 and 9 in which T cell velocity increases, yet the cells remain mostly confined [9]. We have interpreted this behavior as a switch to a rapid surveillance mode in which the T cells vigorously search their local environment for antigen bearing or infected cells [9]. Blocking OVA peptide-MHC complexes with the 25D.1 mAb at day 7 recapitulated the abrupt increase in cell velocities, suggesting that the presence of antigen bearing cells is partly responsible for the cell arrest observed [9], presumably because the T cells are engaging these cells. However, it is possible that antigen is only one of several signals that change as the virus is cleared and antigen becomes much more limiting. The production of various inflammatory cytokines and chemokines likely decrease, and other pro-resolving factors increase [14].

In the present work, we further interrogated the roles of antigen and G-protein coupled chemokine receptors (GPCR) in regulating CD8 T cell motility behavior. Strategies to block signals from peptide-MHC and GPCR signaling were employed. We also artificially re-introduced antigen in the form of soluble peptides to see if this reversed the rapid surveillance and increased cell arrest. We found that both the presence of antigen and GPCR signals (presumably chemokine driven) regulated CD8 T cell motility, reinforcing the conclusions that these are major signals regulating the T cells in the tissue.

## Materials and Methods

### Mice

Male C57BL/6 (B6) mice were obtained from Jackson Laboratories (Bar Harbor, ME) and primary inoculation with influenza virus was performed between 8 and 12 weeks of age. All animals were housed in the University of Rochester Vivarium facilities under specific pathogen-free conditions using microisolator technology. All animal experiments were performed according to the protocol approved by the University Committee for Animal Research (the University’s IACUC).

### Cell transfer and infection

A line of GFP expressing OT-1 animals is maintained and used as donors for adoptive transfer of C57BL/6 naïve animals prior to infection. Whole splenocytes are isolated, and 1×10^6^ splenocytes contained in 200µl of DPBS are transferred via tail vein. 24 - 48 hours post transfer, the animals are sedated with 120 mg/kg of Avertin. Once animals are sedated and exhibit no pedal reflex, 30 µl of X31-OVA-1 (2.7×10^3^ EID50/ml) is administered intra-nasally. Animals are monitored until fully recovered from anesthesia, and then daily for weight-loss.

### In vivo Multiphoton Microscopy (IVMPM)

All images were collected by an Olympus FVMPE-RS system (Olympus, Center Valley, PA) using an Olympus 25× water objective (XLPLN25XWMP2, 1.05NA). The system was equipped with two two-photon lasers: Spectra-Physics InSightX3 (680nm-1300nm, Spectra-Physics, Santa Clara, CA) and Spectra-Physics MaiTai DeepSee Ti:Sapphire laser (690nm-1040nm). There were four Photon Multiplier Tubes (PMTs) and two filter cubes (Blue/Green cube: 420-460nm/495-540nm, Red/Far Red cube: 575-630nm/645-685nm) for multi-color imaging. A galvanometer scanner was used for scanning, and all images were acquired at ~1 frame/s. PMT gains for all imaging were used between 500 and 700 airy units in the Olympus Fluoview software.

### Sedation and Anti-convulsants

Prior to surgery, the animal is anesthetized with pentobarbital (65 mg/kg). During imaging the level of anesthesia is maintained with isofluorane, administered at 0.5-2% as necessary based on heart rate. Pancuronium bromide (0.4mg/kg) is administered prior to imaging, to prevent movement of the imaging area.

### Monoclonal antibodies

The following antibodies were used in the live imaging experiments. Those in this list were injected at 100µg/100µl intra-venous, just prior to live imaging on the multi-photon microscope: LEAF purified Mouse IgG1 BioLegend #400153; and anti-mouse MHC Class I (H-2K^b^) bound to SIINFEKL peptide (OVA residues 257-264) BioXCell #BE0207 [15].

### The introduction of Pertussis toxin

Ptx (List Biological Laboratories #180), was injected at 1.5µg/200µl. This was performed intra-venously 5 hours prior to live imaging by multi-photon microscopy.

### The introduction of OVA siinfekl peptides

To introduce additional antigen into the mice at days 8 and 9 after infection, 100µg siinfekl peptide dissolved in PBS was injected intravenously immediately after sedation and just prior to imaging [16].

### Surgical set-up

The hair is removed with a shaver from one hind leg, thigh to groin, exposing the skin for the MouseOX sensor. The hair is removed from the thoracic area with scissors and/or a shaver. The animal is placed in a supine position, on a warming blanket, once a surgical plane of anesthesia is determined by lack of both pedal and palpebral reflex. The coat is opened from below the chin to the top of the ribcage. The salivary glands are separated to reveal the muscles covering the trachea. Using round forceps, the muscles are separated to expose the trachea. The forceps are then inserted beneath the trachea to lift and separate it from the muscle and mouse body. A small flexible plastic support is placed in the space created by the forceps to permanently hold the trachea above and separated from the muscle, surrounding tissue and coat.

The animal is moved to the previously warmed stage. A small incision is made between the cartilage rings below the larynx. The steel cannula is inserted into the opening in the trachea until it reaches just below the sternum. The cannula provides physical immobilization of the trachea. The cannula is secured on the stage with a support that holds it in position, so it is correctly aligned with the trachea. It is held in place with 2 screws that prevent it from moving in transport, or during the attachment of the respirator. The forepaws are secured to the stage with surgical tape to maintain position of the mouse body on the stage. A few drops of saline are placed on the exposed tracheal tissue to prevent drying.

### Maintaining blood O2 saturation and temperature

The cannula is quickly attached to a Harvard Inspira Advanced Safety Ventilator, and both 100% O2 and 0.5% isofluorane flow is started, according to mouse weight. The MouseOX Plus thigh sensor is attached to the exposed thigh and monitoring is begun immediately. Oxygenation levels are maintained at 95% and heart rate ranges between 250 and 600 beats per minute. The rectal body-temperature is continuously monitored and maintained using a small animal temperature controller that is connected to a rectal probe and a feedback-regulated rodent heating pad. After achieving stable physiology, and verification of lack of both pedal and palpebral reflex, the pancuronium bromide (0.4 mg/kg) is administered, based on body weight. The saline covering the exposed trachea is blotted away and replaced with 0.05% agarose to seal the exposed area. Once the agarose is solidified, a support ring is placed over the imaging area and covered with a piece of plastic wrap. Approximately 5 ml of water is pooled over the imaging area to submerse the objective.

### Humane end point

Imaging is performed with the animals fully sedated as described above. At the end of an imaging session, which last one to two hours, all animals are euthanized by increasing isofluorane dose and ceasing respiratory support. The animal’s heartbeat is continually measured, and the animal is considered euthanized when the heart stops beating. This occurs within one to five minutes after imaging has stopped. The numbers of animals used are indicated in the figure legends for each set of data.

### Data Analysis

The imaging data was analyzed using Volocity 6.3 (Perkin Elmer, Waltham, MA) and Imaris 9.3 (Oxford Instruments, Concord, MA) to determine motility parameters. Each cell track was manually checked and ambiguous tracks, such as two cells coming together and indistinguishably separating or cells with partial tracks caused by leaving the imaging field, were removed from the analysis.

## Results

### Pertussis toxin sensitivity

Chemokines and chemokine receptors are critical in directing T cells to sites of infection. CXCL9 and CXCL10 from epithelial cells are strongly induced by influenza infection [14, 17]. The receptor for these chemokines, CXCR3, is expressed on activated CD8 T cells and is required for efficient infiltration of the infected respiratory tract [18]. We therefore reasoned that interfering with chemokine signaling would affect T cell infiltration into the infected airways. Pertussis toxin (Ptx) interferes with GPCR signaling which is the main pathway for most chemokines including CXCL9 and CXCL10 [19]. Naïve OT-I CD8 T cells were adoptively transferred one day prior to intranasal inoculation with recombinant X31-OVA (SIINFEKL) virus. The animals were then treated on days 6-9 with Ptx five hours prior to imaging [15]. Ptx sensitivity varied over the course of infection. On days 6 and 7, Ptx treatment reduced cell velocities, displacement, and increased confinement (defined by the meandering index) (Figure 1A-C; Supplemental Figure 1A-C). However, on day 8 when T cell velocities naturally reach a nadir and confinement is high (Supplemental Movie S1), the opposite effects were observed. T cell velocities increased with Ptx, and confinement decreased (Figure 1A-C; Supplemental Figure 1A-C). On day 9 when cell velocities normally increase markedly, but transiently [9] (Supplemental Movie S1), the effects of Ptx slowed the cells and decreased confinement, with the largest fold-changes observed. Overall, the effects of Ptx-mediated inhibition of signaling through GPCRs varies over the course of acute infection. We interpret these results to indicate chemokine signals are needed as the T cells infiltrate the tissue from the bloodstream on days 6 and 7, but by day 8 the majority of cells are arrested (velocities <2 µm/min) and more confined in the control mice, presumably as they have reached the epithelium. For example, the highest number of CD8 T cells obtained by bronchoalveolar lavage occurs on day 8 [20]. This may be explained by the physical environment of the epithelium, potential to arrest in the presence of high chemokine concentrations [21], and/or engagement with antigen bearing cells (ABC), which will be investigated later in this paper. Thus, the effects of Ptx treatment are dynamic, varying with the day after infection as the T cell infiltration and clearance of virus progresses with each day. GPCR signals promote cell infiltration early in the responses but can also cause arrest in areas of the tissue with high concentrations of signals [21]. The significant increases in apparent cell velocities with high confinement that occur on day 9 also appear GPCR dependent, though we cannot rule out other signals that change and contribute to the rapid the movement.

**Figure 1:**
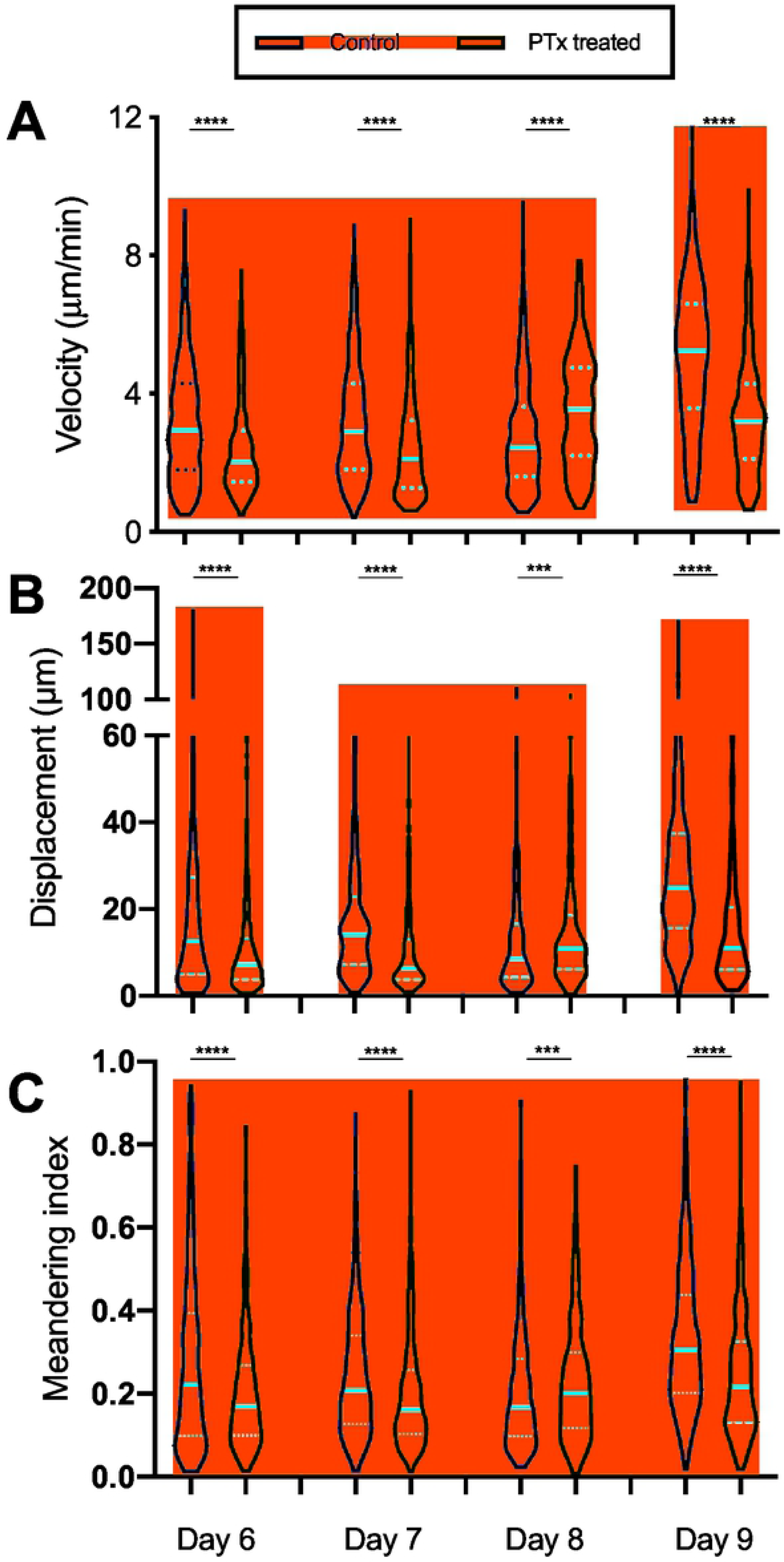
Pertussis toxin treatment effects vary depending on the day of infection. OT-IGFP CD8 T cells were adoptively transferred into naïve hosts one day prior to infection with influenza virus expressing the siinfekl peptide. CD8 T cell motility was tracked by IVMPM and the data analyzed using Volocity software. Ptx was administered intravenously 5h prior to imaging. The motility parameters for velocity (A), Displacement (B), and Meandering Index (C) are plotted as violin plots. The median and standard deviations appear in red. A non-parametric Mann-Whitney test was applied to each pair of data sets for each day of infection. *** p < 0.01; **** p < 0.0001. N = 5 animals per time point

### Distance from the airway

Influenza mostly infects the epithelial cells lining the respiratory tract. If the T cells are responding to chemokine signals that direct them towards the infected epithelium on days 6-8, then we would predict that interference with these signals would result in delayed migration to the airway surfaces. We therefore developed a computational approach to measure the depth of the T cells in the trachea relative to the airway surface. The measurements were conducted from days 6-9 of infection. In the control mice, T cell abundance closest to the airways peaked on day 7 and actually decreased on days 8 and 9 (Figure 2). Consistent with the predictions, Ptx treated mice demonstrated delayed accumulation of T cells close to the airway surfaces that was particularly evident on days 6-7 (Fig. 2). By day 8 the proportion of T cells closest to the airways had eventually caught up to some extent in the Ptx treated mice, but never reached the peak levels observed in the controls (Fig. 2). These results indicate that the T cells rely on chemokine signals for optimal migration to the site of infection in the epithelium.

**Figure 2:**
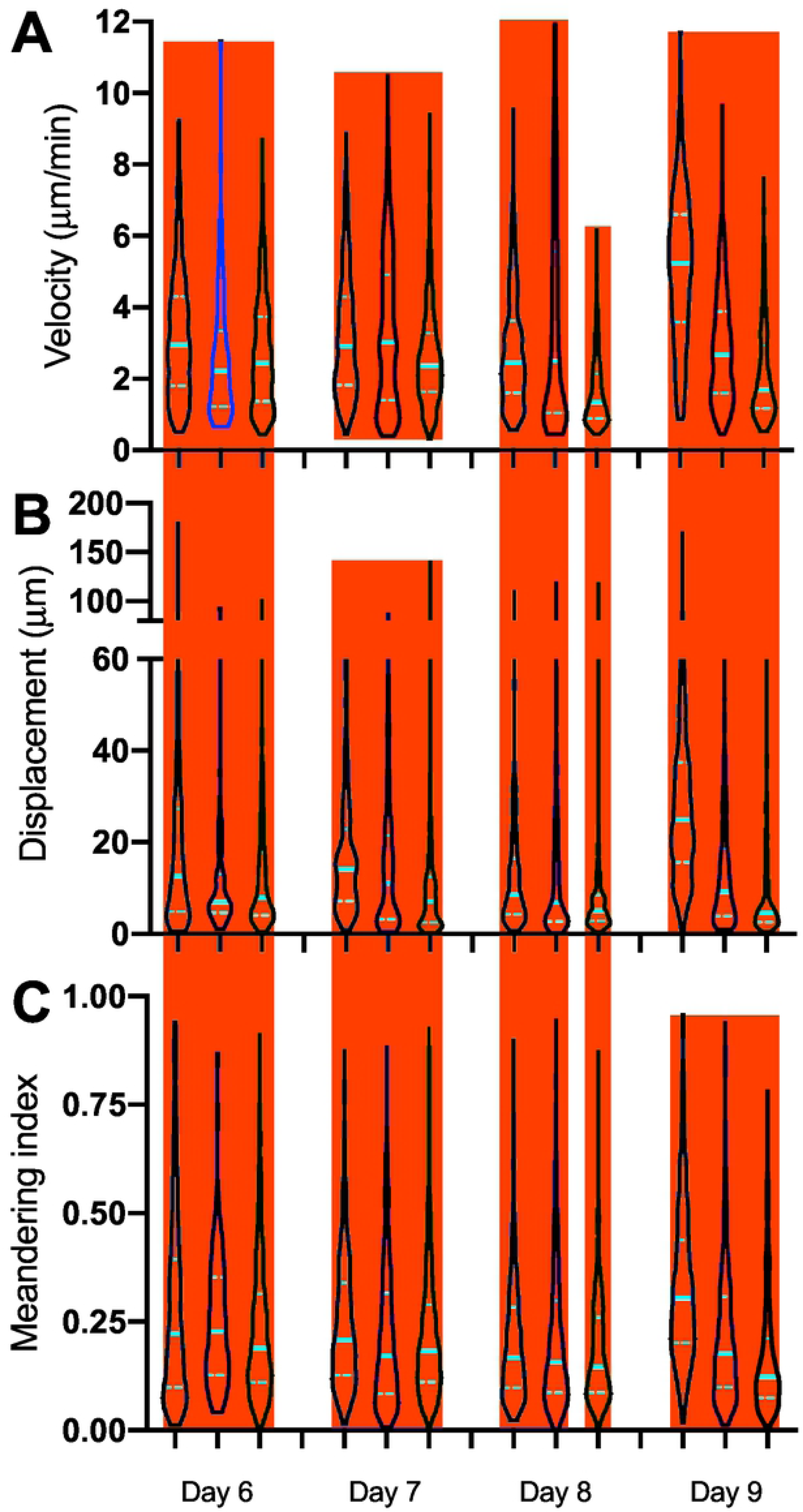
Penetration depth of OT-I-GFP CD8 T cells with and without pertussis toxin treatment. The data from each day of IVMPM in influenza infected mice was processed by a custom module developed for Imaris software. (A) Tissue depth is measured relative to the Second Harmonic Signal (SHG) from collagen in the outer sheath to the airway lumen which is easily demarcated in the imaging by a charge in light diffraction at the tissue-air interface. (B) The histograms indicate the number of cells at any given depth. Units are relative and set to an arbitrary scale. The median depth and total number of cellular events is indicated by the numerical values at the bottom of each plot. The differences in penetration depths were significantly different (p < 0.0001) between controls and Ptx treated pairs of data for each day using non-parametric Mann-Whitney tests. N = 5 animals per time point.

### Peptide-MHC sensitivity

It was not possible in our studies to unambiguously distinguish infected cells and antigen cross-presenting cells, so we are using the term antigen bearing cells “ABC”. We hypothesized that chemokine signals and the presence of ABC that can engage T cells in a peptide-MHC specific manner cooperate to retain effector CD8 T cells at the epithelium where the infection is present. We expected encounter with ABC to result in cell arrest, at least transiently, as the CD8 T cells kill their targets or are further activated by APC. To investigate the role of antigen encounter, as above, naïve OT-I CD8 T cells were adoptively transferred and the mice infected the next day with X31-OVA. To inhibit peptide-MHC (pMHC) specific T cell interactions, we used a monoclonal antibody (25-D1) generated against the SIINFEKL/H2Kb class I MHC-peptide complex to treat the animals immediately prior to imaging [15]. There were no significant effects on day 6, an early time point when the number of T cells engaged with ABC in the tissue is expected to be relatively low. However, on days 7-9 specific pMHC blocking unexpectedly and significantly decreased cell velocities and displacement (Figure 3A, B) compared to untreated and isotype treated controls but had little effect on the confinement indices (Figure 3C). For unknown reasons, the isotype control affected cell motility parameters on day 9, but not on other days. The anti-pMHC treatment still further reduced all parameters on day 9 compared to either the isotype or untreated controls. These results suggest TCR engagement with pMHC has a role governing the motility behavior of the T cells, possibly by further activating the cells in the tissue.

**Figure 3:**
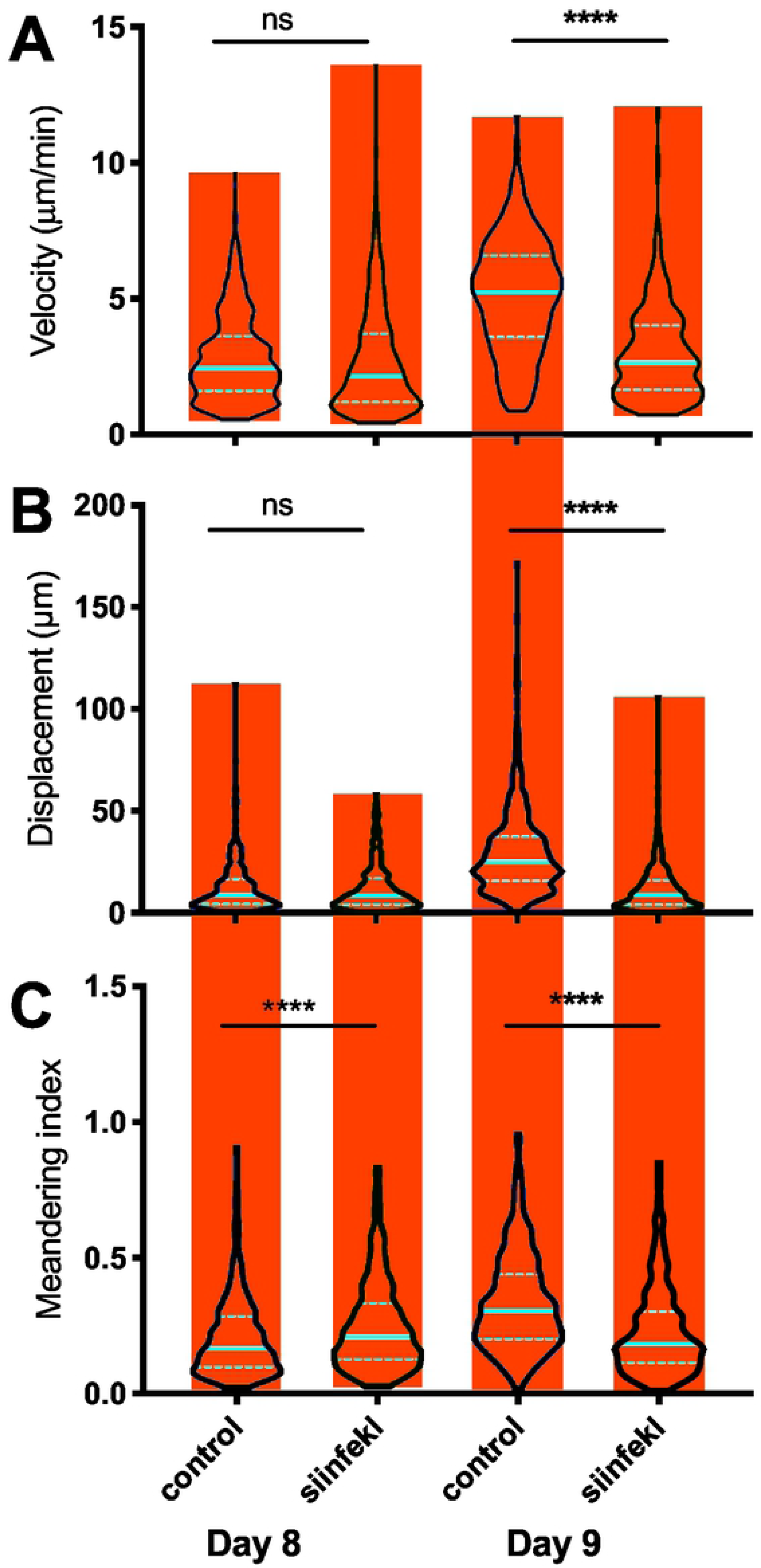
Effects of specific peptide-MHC mAb blocking on CD8 T cell motility during acute influenza infection. OT-I-GFP T-CD8 T cells were adoptively transferred into naïve hosts one day prior to infection with influenza virus expressing the siinfekl peptide. CD8 T cell motility was tracked by IVMPM and the data analyzed using Volocity software. 100µg of OVA pMHC mAb 25-D1 or a control isotype matched IgG were administered just prior to imaging by an intravenous route. The motility parameters for Velocity (A), Displacement (B), and Meandering Index (C) are plotted as violin plots. The median and standard deviations appear in red. A non-parametric Mann-Whitney test was applied to each pair of data sets for each day of infection. ns = not significant; **** p < 0.0001. N = 5 animals per time point.

### Re-introduction of antigen

We reasoned that the changes on day 9 were the result of a significant reduction in signals received from ABC as the virus is cleared [9]. To address this, we re-introduced antigen in the form of soluble SIINFEKL peptides on days 8 and 9. The introduction of peptides had no effect on motility parameters on day 8 when virus and, presumably, ABC are still abundant. However, on day 9, the introduction of peptides resulted in significantly decreased velocities and displacement, increased numbers of cells arrested, and heightened confinement compared to vehicle controls (Figure 4A-C). This is consistent with the results of pMHC blocking on day 8, and strongly suggests that the balance of antigen signals in the presence of continued chemokine signaling result marked changes in velocity and confinement that occur between day 8 and day 9. This is consistent with the interpretation that the still highly activated T cells switch to a rapid surveillance of the surrounding cells for the presence of virus or residual antigen and ABC.

**Figure 4:**
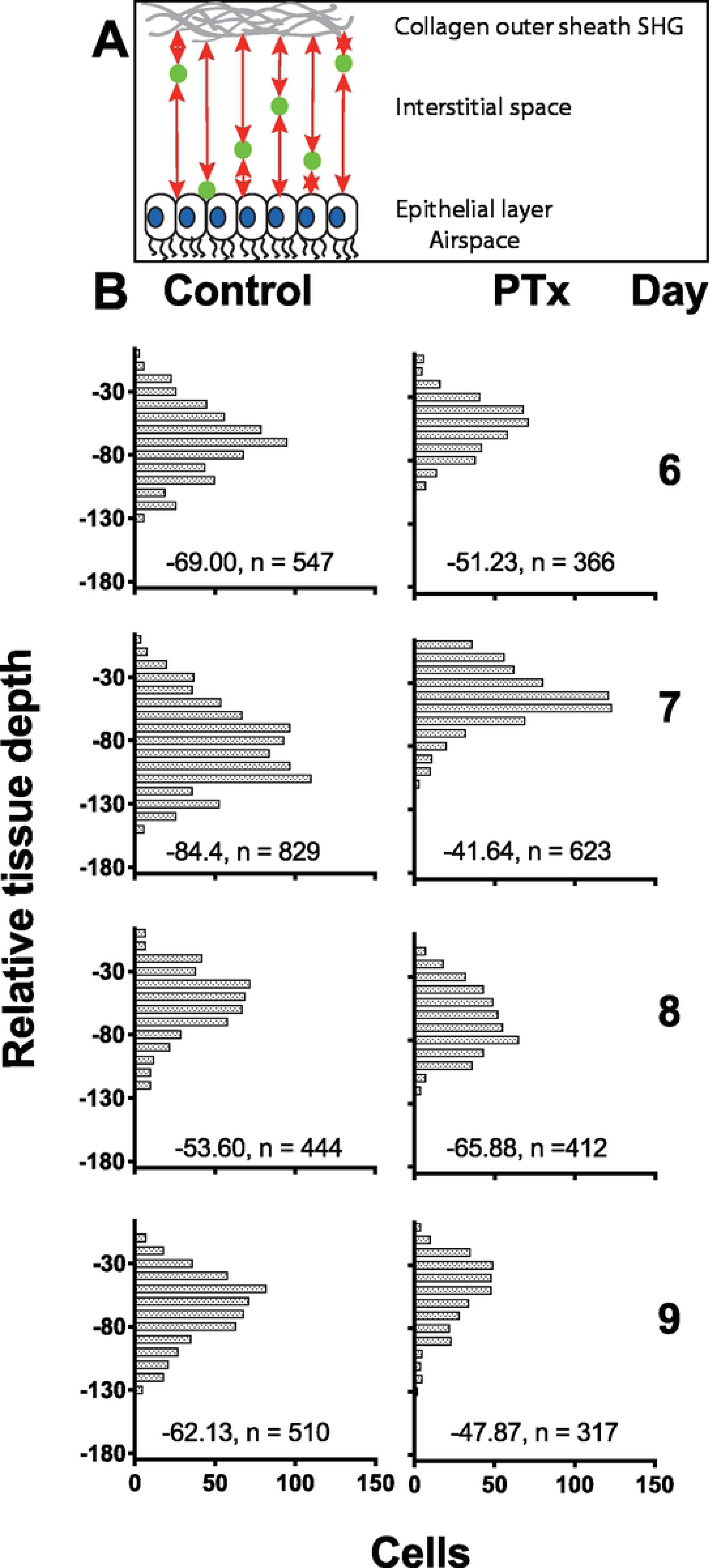
Administration of exogenous OVA siinfekl peptide restores T cell arrest on day 9. Host animals received adoptively transferred CD8 T cells and were infected with influenza as described in Material and Methods. 100μg OVA siinfekl peptides in saline were injected intravenously during the imaging. CD8 T cell motility was tracked by IVMPM and the data analyzed using Volocity software. The motility parameters for velocity (A), Displacement (B), and Meandering Index (C) are plotted as violin plots. The median and standard deviations appear in red. A non-parametric Mann-Whitney test was applied to each pair of data sets for each day of infection. ns = not significant; **** p < 0.0001. N = 5 animals per time point.

## Discussion

Chemokines and chemokine receptors play major roles in guiding T cell migration into the infected airways. During influenza infection, secretion of CXCL9 and CXCL10 from the inflamed epithelium recruits CD8 T cells expressing CXCR3 chemokine receptors [18]. We have also shown that neutrophil-derived CXCL12 is an important enhancer of CD8 T cell infiltration into the infected airways, signaling via CXCR4 [22]. Neutrophil depletion delays CD8 T cell recruitment into the tissue as well as impairing virus clearance [22, 23]. We observed that inhibition of GPCR signals using Ptx treatment significantly reduced virus-specific CD8 T cell motility on days 7 and 8 when T cell infiltration is at its height, reducing cell velocities, decreasing displacement, and increasing measurements of T cell confinement (meandering index). This suggests chemokine signals are critical for efficient tissue infiltration, as Ptx treatment led to slower penetration of CD8 T cells to the epithelial surface. Individual T cells are known to migrate using a random walk [24]. This is consistent with the idea that chemokine signals promote chemokinesis and directional accumulation at the infected epithelium. Interestingly, at the peak of the T cell response on day 8 when cell numbers are highest and virus is still present, Ptx treatment relieved cell arrest and confinement. This is consistent with the idea that once the majority of T cells have reached the infected epithelium, they arrest through a combination of physical confinement (the epithelium is narrow in cross section, and there is dense extracellular matrix underlying the epithelial surface), high CXCL9 and CXCL10 chemokine concentrations at the epithelium, which can contribute to arrest, and presumably engagement with antigen bearing cells in the form of either APC or infected targets. This is discussed further below.

On day 9 when virus is cleared, the CD8 T cells normally exhibit an increase in cell velocities, and yet remain confined. Imaging of this behavior shows rapid extension and retraction of T cell protrusions, but little distance traveled (compare Supplemental Movies S1 and S2). We have interpreted this collective behavior as rapid surveillance for residual antigen and virus by the highly activated effector T cells. Treatment with Ptx at this time point significantly decreased all the motility parameters, suggesting a requirement for GPCR signals for optimal surveillance of the tissue as the infection resolves. How chemokines and chemokine receptors affect changes to T cell motility on day 9 is unresolved. The chemokines CXCL9 and CXCL10 are expressed constitutively by airway epithelium, and increase during infection [25, 26]. In addition to potentially providing chemotactic and chemokinetic signals, they may be needed for integrin activation [27], each of which could contribute to the cell behavior we observe, though we have not formally tested these mechanisms in our model.

The experiments blocking MHC with specific monoclonal antibodies showed significant effects on CD8 T cell motility. However, the effects were not entirely as expected. If we take the hypothesis that engaging antigen bearing cells will cause cells to arrest, at least briefly, as has been observed in other T cell motility studies [28], then we would predict that blocking antigen signals would partially or completely reverse cell arrest. On day 6 there were no differences in the motility parameters between pMHC mAb treated mice and isotype control mice, likely because few T cells had found their targets. But on days 7-9, velocities were consistently reduced with pMHC mAb treatment, with greatest magnitude of the effect on day 8, a time when we expect maximal engagement with ABC. There was still an effect on day 9, after virus is cleared and no productively infected cells are detectable, suggesting there may be residual antigen. Confinement indices were not affected at most time points, suggesting chemokine and/or physical factors maintain confinement and arrest. We conclude that effector CD8 T cells in the airways need T cell receptor signals that stimulate optimal motility.

Supporting the idea that antigen signals are limiting on day 9, resulting in the observed increased motility as the cells search for residual antigen, the addition of soluble siinfekl peptides into the mice resulted in significantly decreased velocities and increased confinement. There was little effect of the peptides added on day 8. These results further support the notion that signals through the TCR affect both movement and arrest depending on the day of infection.

Overall, the combined results of Ptx treatment, pMHC blockade, and re-introduction of exogenous antigen paint a consistent picture of how chemokine signals and T cell engagement with ABC each independently regulate CD8 T cell motility in the acute phase of the infection. The individual effects vary as the infection and infiltration of the T cells into the tissue progress. It is important to consider that T cell responses in the tissues are dynamic and evolve as the immune response begins, reaches a peak, and then resolves. Studies of T cell motility that use a single time point may not reveal the full picture of how different signals regulate T cell migration over the course of an infection.

---

**Supplemental Movie S1: Movie S1:** Days 8 and 9 after infection sequentially showing IVMPM of OT-I-GFP+ CD8 T cells moving in the trachea of X31-OVA-I influenza infected mice. The day 8 movie depicts both arrested and motile cells in the field. Image traverses the entire depth of the trachea. N = 1 mouse. The day 9 movie depicts many cells exhibiting a rapid “scanning” motility behavior with low displacement. Image traverses the entire depth of the trachea. N = 1 mouse.

**Supplementary Figure S1: Pertussis toxin treatment effects vary depending on the day of infection.** OT-I-GFP CD8 T cells were adoptively transferred into naïve hosts one day prior to infection with influenza virus expressing the siinfekl peptide. CD8 T cell motility was tracked by IVMPM and the data analyzed using Volocity software. Ptx was administered intravenously 5h prior to imaging. The individual cell motility parameters for velocity (A), Displacement (B), and Meandering Index (C) are plotted using symbols. The median and standard deviations appear in red. A non-parametric Mann-Whitney test was applied to each pair of data sets for each day of infection. *** p < 0.01; **** p < 0.0001. N = 5 animals per time point.

**Supplementary Figure S2: Effects of specific peptide-MHC mAb blocking on CD8 T cell motility during acute influenza infection.** OT-I-GFP T-CD8 T cells were adoptively transferred into naïve hosts one day prior to infection with influenza virus expressing the siinfekl peptide. CD8 T cell motility was tracked by IVMPM and the data analyzed using Volocity software. 100μg of OVA pMHC mAb 25-D1 or a control isotype matched IgG were administered just prior to imaging by an intravenous route. The individual cell motility parameters for Velocity (A), Displacement (B), and Meandering Index (C) are plotted using symbols. The median and standard deviations appear in red. A non-parametric Mann-Whitney test was applied to each pair of data sets for each day of infection. ns = not significant; **** p < 0.0001. N = 5 animals per time point.

**Supplementary Figure S3: Administration of exogenous OVA siinfekl peptide restores T cell arrest on day 9.** Host animals received adoptively transferred CD8 T cells and were infected with influenza as described in Material and Methods. 100μg OVA siinfekl peptides in saline were injected intravenously during the imaging. CD8 T cell motility was tracked by IVMPM and the data analyzed using Volocity software. The individual cell motility parameters for velocity (A), Displacement (B), and Meandering Index (C) are plotted using symbols. The median and standard deviations appear in red. A non-parametric Mann-Whitney test was applied to each pair of data sets for each day of infection. ns = not significant; **** p < 0.0001.

## Acknowledgements

We thank Dr. Yurong Gao of the Multiphoton Core Lab for her technical support. This work was funded by NIH/NIAID/DAIT grant number P01-AI102851 and MS was supported by National Institute of General Medical Sciences Medical Scientist Training Program T32 GM007356.

